# Towards deciphering the structure of long homozygous stretches in cattle genome

**DOI:** 10.1101/2021.03.31.437853

**Authors:** Michael Grigorievich Smaragdov

## Abstract

Up-to-day there is no as universally accepted software tool and threshold parameters to identify runs of homozygosity (**ROH**). The relative position of POH segments in the cattle genome has not been studied extensively. Specific objective of this study was to evaluate the effect of allowed missing and heterozygous SNPs in ROH on their number, on the estimate of inbreeding level, and on structure of ROH segments in the cattle genome. In this study 371 Holsteinized cows from six herds were genotyped with BovineSNP50 array. To identify ROH, the consecutive and sliding runs were carried out with detectRUNS and Plink tools. Neither effect was shown for missing SNPs genotype calls. Allowing even one heterozygous SNP resulted in significant bias of ROH data. Furthermore, the sliding runs identified less ROH than consecutive runs. The mean coefficient of inbreeding across herds was 0.111 ± 0.003 and 0.104 ± 0.003 based on consecutive and sliding runs respectively. It was shown how, using the heterozygous SNPs in ROH, may be possible to derive a distribution of ROH segments in the cow genome. We suggested it was similar to normal distribution. Furthermore, frequency of ROH in the chromosomes did not depend on their length. Of 29 chromosomes, the most abundant with ROH were BTA 14, BTA 7, and BTA 18. The result of this study confirmed more accurately identification of ROH with consecutive runs, uneven their distribution in the cattle genome, significant bias of the data due to allowing heterozygous SNPs in ROH.

## Background

Inbreeding in dairy cattle is an inevitable phenomenon of artificial selection. Traditionally coefficient of inbreeding is calculated based on ancestry [1]. With the advent of SNPs arrays become possible to investigate autozygosity at a previously unattainable level [2]. In fact, due to runs of homozygosity (**ROH**) approach the analysis of the animal genome for extended homozygous SNPs stretches is still ongoing. The primary cause of autozygosity in livestock measured with ROH is assumed to be inbreeding [2] or consanguineous marriage in humans [3]. For identifying of ROH, the softwares based on either identity by descent (IBD) GERMLINE [4], or Hidden Markov Model (HMM) Beagle [5] and BCFtools [6] were elaborated. An addition, the software based on scanning with SNPs window Plink [7], overlapping sliding window SNP101[8], or both consecutive and sliding runs detectRUNS [9] as well software based on other scripts [10,11], cgaTON [12] have been provided. There is the commercial software SNP & Vriation Suite (Golden Helix SNP & Variation Suite). It was shown that software based on HMM and IBD occurred to be inferior to other software mentioned above [13].The major difficulty faced by scientists is the lack of consistent criteria among studies regarding the threshold values in each parameters analyzed to define the ROH [2]. The most crucial parameters to be used in software are the number of allowed heterozygous or missing SNPs calls in ROH. There is inconsistency among thresholds to be applied in published studies. Some authors disallowed any number of heterozygous SNPs in ROH [14–17], other allowed one, two and more heterozygous SNPs depending on the length of ROH segments [18–25]. Anyway, Ferencakovic et al. [22] suggested that allowing some number of genotype errors in long ROH may minimize underestimation of those segments. Though, Mastrangelo et al. [24] showed small increasing in inbreeding coefficient, if heterozygous genotypes were allowed.

There are relatively a few studies assessing what a set of these parameters is optimal to detect ROH, so as to better understand their effect on identified autozygosity. Ferencakovic et al. [22] have shown that SNP array density and genotyping errors introduce patterns of bias in the estimation of autozygosity. These authors observed that allowance of the heterozygous SNPs in ROH may has led to merging of adjacent ROH segments that resulted in biased estimates of the ROH number. Based on simulation data Howrigan et al. [13] recommended not allowing any heterozygous SNPs existence in a called ROH. Summarizing, there is currently no generally accepted opinion about the reasonable number of heterozygous SNPs in ROH to avoid bias in the ROH data.

The Leningrad region is the highest average milk yield producing region in Russia, with approximately 60,000 cows of Holsteinized Black and White cattle. Dutch, Danish, and Swedish Black and White bulls and heifers were imported to Russia during the 1930s. The Black and White breed was officially registered in Russia in 1959. To improve milk traits of Black and White cattle, local farmers started to use imported from USA (since 1978) and The Netherlands (since 2002) Holstein bulls and semen. Currently, the commercial Russian Black and White cattle population can be considered as Holstein due to long-term crossing only with Holstein bulls.

Main objectives of this study were: (i) to estimate the number of ROH segments in the cow genome with consecutive and sliding runs, (ii) to assess their dependence on allowed number of heterozygous and missing SNPs calls in ROH, (iii) to quantify the coefficient of inbreeding of the cows in the herds studied, (iv) to compare the results obtained by these two approaches (v) to elucidate the structure and distribution of ROH in cow genome.

## Materials and Methods

### Ethics statement

Th*is* study does not involve any endangered or protected species. Samples were derived from the Committee on Agro-Industrial Complex of Leningrad region. The permission to carry out the sampling at each farm was obtained directly from the owners. All the samples were collected during routine veterinary checks and in accordance with local/national regulations and ethical rules in force at the time of sampling. The study was approved by the Ethical & Animal care committee of the RRIFAGB — Branch of the L. K. Ernst Federal Science Centre for Animal Husbandry. Animals were handled with respect to the Russian Federal Law No. 498-FZ on Responsible Treatment of Animals and on Amendments to Certain Legislative Acts of the Russian Federation.

### Animal resources and SNPs genotyping

Data and genotypes were obtained from Committee on Agro-Industrial Complex of Leningrad region. Cows genotypes were available from six breeding herds located in Leningrad region (Russia) born in range from 2010 to 2013. Animals for genotyping were selected by farmers with regarding the pedigree structure of the herd. Number of animals genotyped depends on number of sires used in herd at least one daughter per sire. In case of multiple daughters per single sire, the sires of dams were different. Sampled animals included 8–15% of total number of milking cows in herds. Altogether, 371 cows were genotyped with BovineSNP50 v.2.0 array (Illumina Ca. USA). In the first quality control step, SNPs with quality score less than 0.7 were removed. In the second step, genotyping quality control was done with PLINK 1.9 [7]. Only autosomal chromosomes were considered. In data set missing rate per SNP was no more than 5% and probability of deviation from Hardy-Weinberg equilibrium (HWE) was less than 1.0E-03. SNPs with MAF < 0.01 were removed resulted in 48,108 SNPs. Total genotyping rate was > 0.99.

### Detection the runs of homozygosity

The ROH segments were identified using detectRUNS [9] implemented in R environment [26], and Plink tool [7]. The parameters and threshold applied to define ROH with detectRUNS for consecutive runs method were: (i) minimal number of SNPs needed to define segment as a ROH: 15 and 20, (ii) number of missing calls allowed in a ROH segment: 0 - 4, (iii) number of heterozygous calls allowed in a ROH segment: 0 - 240, (iv) minimal length of ROH segments: 250 Kb, (v) maximal gap between ROH segments: 1 Mb.

For sliding window method the parameters and thresholds were: (i) window size: 15 SNPs and 20 SNPs, (ii) threshold: 0.05, (iii) minimal number of SNPs needed to define segment as a ROH: 15 and 20, (iv) number of missing calls allowed in a ROH segment: 0 - 4, (v) number of heterozygous calls allowed in a ROH segment: 0 - 16, (vi) minimal length of ROH: 250 Kb, (vii) maximal gap between ROH segments: 1 Mb, (viii) minimal allowed density of SNPs: 1 SNP per 1 Mb.

The parameters and threshold applied to define ROH with Plink were: (i) sliding window: 20 SNPs, (ii) the proportion of homozygous overlapping windows: 0.05, (iii) the minimum number of SNPs in ROH: 20, (iv) density was one SNP per 50 Kb, (v) the number of missing SNPs: zero, (vi) the number of heterozygous SNPs: zero.

Inbreeding coefficient was calculated as the sum of ROH length of an animal divided by the total length of the autosomes covered by SNPs (2508.706681 Mb). A probability (quantile-quantile (Q-Q) plot was performed in R environment [26].

## Results

In this section the effect of the number of heterozygous and missing SNPs on ROH data was evaluated by consecutive and sliding runs. Primarily, the effect of missing SNPs allowed in ROH on the data was evaluated by consecutive and sliding runs. Both methods were not show any effect on ROH data when from one to four missing SNPs calls were allowed in ROH. Therefore, to evaluate the ROH findings further, this value has been fixed to zero.

### Effect of heterozygous SNPs on ROH data

#### ROH identification based on consecutive runs

To detect the number of ROH segments in the cow genome, 15 SNPs (S1 Table) and 20 SNPs (Table 1) consecutive runs across cows genome were carried out. When ROH segments were not interrupted with heterozygous SNPs the mean number of ROH was greater in 1.9 times at 15 SNPs runs. In fact, the average number of ROH across herds was 182.0 ± 2.8 at 15 SNPs runs, as compared with 94.4 ± 2.7 at 20 SNPs runs. To avoid an autozygous ROH overestimation due to short ROH segments, the 20 SNPs runs were used further.

**Table 1.** Estimated mean (± SE) ROH number in the herds based on 20 SNPs consecutive runs

| Herd | 1 | 2 | 3 | 4 | 5 | 6 |
| --- | --- | --- | --- | --- | --- | --- |
| 0* |  |  |  |  |  |  |
| The mean number of ROH | $99.6 \pm 6.5$ | $90.5 \pm 1.0$ | $91.5 \pm 1.4$ | $106.4 \pm 14.7$ | $93.0 \pm 1.3$ | $91.4 \pm 1.5$ |
| Maximum | 360 | 112 | 148 | 757 | 111 | 125 |
| Minimum | 2 | 66 | 65 | 48 | 75 | 73 |
| 1 |  |  |  |  |  |  |
| The mean number of ROH | $161.9 \pm 10.2$ | $145.4 \pm 1.3$ | $146.5 \pm 1.6$ | $155.8 \pm 12.1$ | $149.2 \pm 1.5$ | $148.8 \pm 1.6$ |
| Maximum | 565 | 179 | 195 | 692 | 175 | 174 |
| Minimum | 7 | 117 | 109 | 82 | 125 | 126 |
| 2 |  |  |  |  |  |  |
| The mean number of ROH | $262.2 \pm 13.1$ | $243.1 \pm 1.6$ | $244.7 \pm 1.8$ | $253.7 \pm 7.4$ | $245.5 \pm 1.6$ | $248.2 \pm 1.8$ |
| Maximum | 761 | 277 | 281 | 564 | 270 | 283 |
| Minimum | 21 | 211 | 215 | 145 | 214 | 207 |
| 4 |  |  |  |  |  |  |
| The mean number of ROH | $583.8 \pm 13.8$ | $573.6 \pm 2.1$ | $579.3 \pm 2.5$ | $582.7 \pm 7.4$ | $576.3 \pm 2.7$ | $580.3 \pm 3.3$ |
| Maximum | 1017 | 625 | 631 | 655 | 636 | 630 |
| Minimum | 116 | 528 | 536 | 383 | 533 | 525 |
| 8 |  |  |  |  |  |  |
| The mean number of | $1161.4 \pm 9.6$ | $1183.3 \pm 2.9$ | $1194.5 \pm 3.3$ | $1182.2 \pm 22.1$ | $1182.5 \pm 3.3$ | $1192.0 \pm 3.7$ |
| ROH |  |  |  |  |  |  |
| Maximum | 1227 | 1254 | 1271 | 1263 | 1233 | 1239 |
| Minimum | 818 | 1126 | 1125 | 224 | 1129 | 1132 |
| 16 |  |  |  |  |  |  |
| The mean number of ROH | 848.4 ± 11.8 | 857.1 ± 2.1 | 862.5 ± 2.4 | 852.2 ± 16.8 | 852.6 ± 2.5 | 860.3 ± 2.9 |
| Maximum | 1297 | 914 | 898 | 1024 | 887 | 893 |
| Minimum | 531 | 810 | 805 | 132 | 805 | 804 |
| 20 |  |  |  |  |  |  |
| The mean number of ROH | 696.1 ± 9.7 | 703.4 ± 1.7 | 707.7 ± 1.9 | 699.4 ± 13.8 | 699.9 ± 1.9 | 706.3 ± 2.3 |
| Maximum | 1069 | 753 | 738 | 840 | 732 | 735 |
| Minimum | 417 | 668 | 659 | 107 | 664 | 663 |
| 60 |  |  |  |  |  |  |
| The mean number of ROH | 257.2 ± 3.4 | 259.6 ± 0.6 | 261.2 ± 0.7 | 258.9 ± 4.9 | 258.2 ± 0.7 | 260.5 ± 0.8 |
| Maximum | 390 | 275 | 272 | 312 | 260 | 272 |
| Minimum | 160 | 248 | 242 | 49 | 248 | 244 |
| 100 |  |  |  |  |  |  |
| The mean number of ROH | 163.9 ± 2.1 | 165.5 ± 0.4 | 166.3 ± 0.5 | 164.6 ± 3.0 | 164.5 ± 0.5 | 165.8 ± 0.5 |
| Maximum | 247 | 174 | 176 | 195 | 173 | 173 |
| Minimum | 103 | 156 | 157 | 38 | 155 | 157 |
| 240 |  |  |  |  |  |  |
| The mean number of ROH | 80.9 ± 0.9 | 81.9 ± 0.2 | 82.2 ± 0.2 | 81.3 ± 1.0 | 81.5 ± 0.3 | 82.1 ± 0.3 |
| Maximum | 114 | 87 | 86 | 93 | 87 | 86 |
| Minimum | 55 | 77 | 78 | 36 | 77 | 77 |

Note: \* - The number of hetrozygous SNPs in ROH

The effect of allowed heterozygous SNPs on the number of ROH segments was evaluated when their values varied from 0 to 240. Descriptive statistics of the data are given in Table 1. The mean number of ROH varied across herds. Nevertheless, between herds mean values were insignificant when heterozygous SNPs were disallowed. The large variation of the ROH number among the cows in fourth herd was observed. This result was due to large number of ROH at one cow (757 ROH segments). Exclusion this cow resulted in the mean ROH number for forth herd 85.9 ± 2.1. As a result, this herd, using t-test estimated with Bonferroni correction, became significantly differed from fifth herd (P < 0.05). Initially, the average number of ROH increased by more than 10 times from 94.4 ± 2.7 up to 1182.7 ± 4.9 when heterozygous SNPs in ROH were allowed from 1 to 8 (Table 1). Then, the mean value decreased to 81.7 ± 0.2 when the number of heterozygous SNPs was increased to 240.

The length of ROH segments was classified into five categories (1-2 Mb, 2-4 Mb, 4-8 Mb, 8-16 Mb, and >16 Mb) (Table 2). The most abundant in the number of ROH was 1-2 Mb class. Significant increase in the number of ROH in this class occurred when 15 SNPs runs were used (Table 2 vs. S2 Table). The largest proportion of ROH number had the same class up to eight allowed heterozygous SNPs, and then its proportion begun noticeable to decline (Table 2). The same type of the distribution was faithful for 2-4 Mb class but with other values of allowed heterozygous SNPs. Namely, the number of ROH began to increase when two heterozygous SNPs were allowed, then the distribution peaked at 16 heterozygous SNPs and gradually decreased further (Fig 1). This type of distribution was the same for 4-8 Mb class but with a lower maximum and at more number of allowed heterozygous SNPs. For 8-16 Mb class increasing of ROH number occurred when sixty heterozygous SNPs were allowed. The mean length of ROH segments for each class is shown in S3 Table. These values did not vary mach, except segments in >16 Mb class with 240 allowed heterozygous SNPs (S3 Table).

**Table 2.** Estimated mean (± SE) across herds ROH number in length classes based on 20 SNPs consecutive run

| Class (Mb) | 0-2 | 2-4 | 4-8 | 6-16 | >16 |
| --- | --- | --- | --- | --- | --- |
| 0* | 3585<br>± 277 | 1155<br>± 9 | 662<br>± 63 | 319<br>± 32 | 113<br>± 17 |
| 1 | 6792<br>± 560 | 1391<br>± 103 | 704<br>± 63 | 344<br>± 34 | 123<br>± 17 |
| 2 | 12382<br>± 1080 | 1840<br>± 142 | 750<br>± 69 | 362<br>± 31 | 131<br>± 18 |
| 4 | 31023<br>± 2857 | 3552<br>± 308 | 874<br>± 76 | 395<br>± 31 | 139<br>± 18 |
| 8 | 59676<br>± 5699 | 11991<br>± 1084 | 1261<br>± 122 | 463<br>± 34 | 161<br>± 17 |
| 16 | 16178<br>± 1570 | 31742<br>± 3010 | 4476<br>± 387 | 612<br>± 65 | 205<br>± 19 |
| 20 | 5049<br>± 486 | 29569<br>± 2816 | 8089<br>± 747.6 | 738<br>± 76 | 229<br>± 20 |
| 60 | 249<br>± 26 | 490<br>± 46 | 4618<br>± 440 | 10076<br>± 953 | 688<br>± 2 |
| 240 | 62<br>± 7 | 119<br>± 14 | 478<br>± 47 | 528<br>± 51 | 3891<br>± 364 |

Note: \* - The number of allowed heterozygous SNPs in the ROH segments

**Figure 1.**
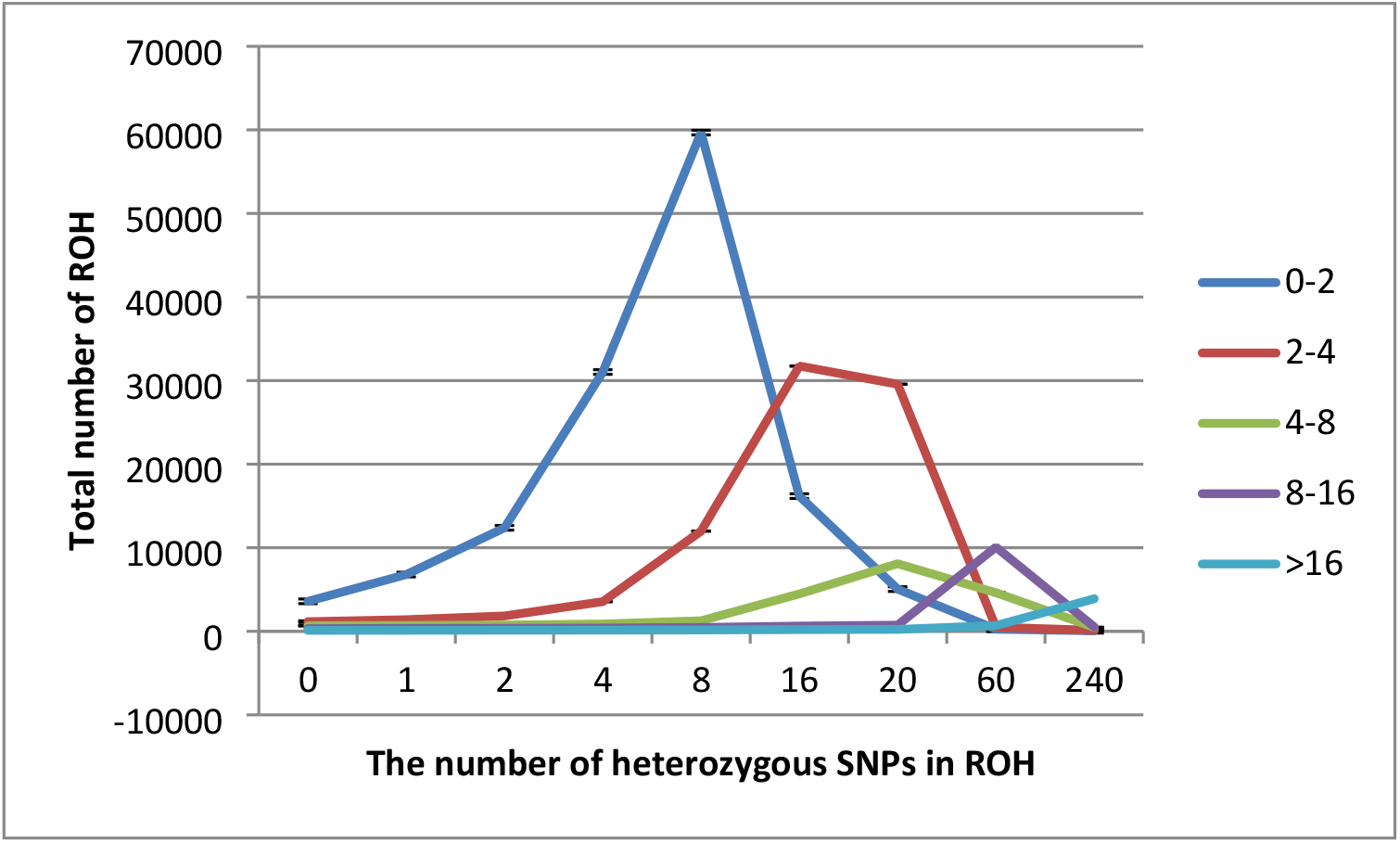
Distribution of ROH in five length classes based on consecutive runs

#### ROH identification based on sliding runs

As for consecutive runs, 15 SNP window (S4 Table) and 20 SNPs window (Table 3) for sliding runs were used. Interestingly, the data for 15 SNPs runs identified both methods nearly coincided (S1 Table *vs*. S4 Table) while for 20 SNPs runs the data differed significantly in 1.9 times (P < 0.001) (Table 1 *vs*. Table 3). Therefore, for this approach as well as consecutive runs, the shorter window was, the shorter ROH segments were identified.

**Table 3.**
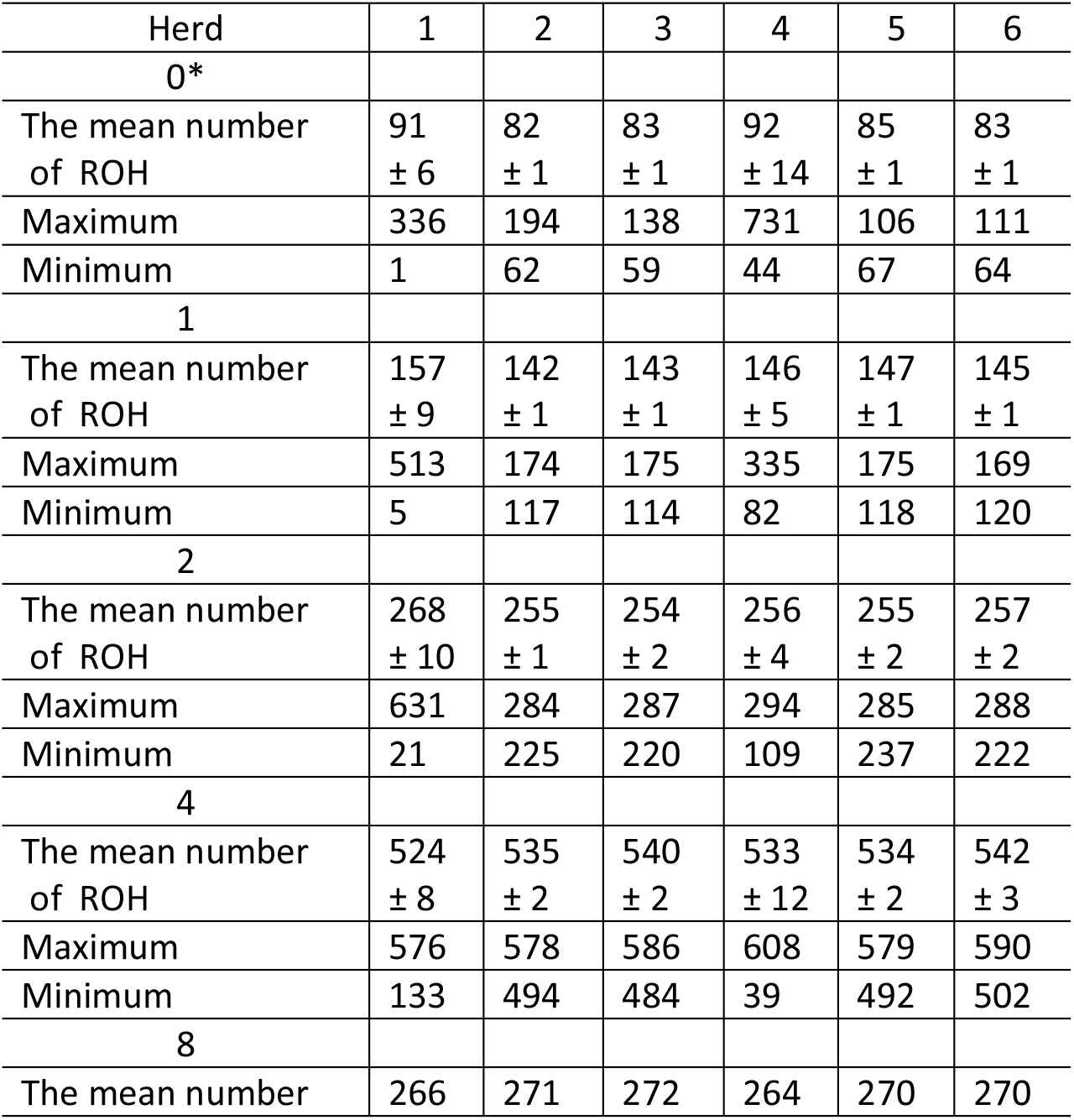
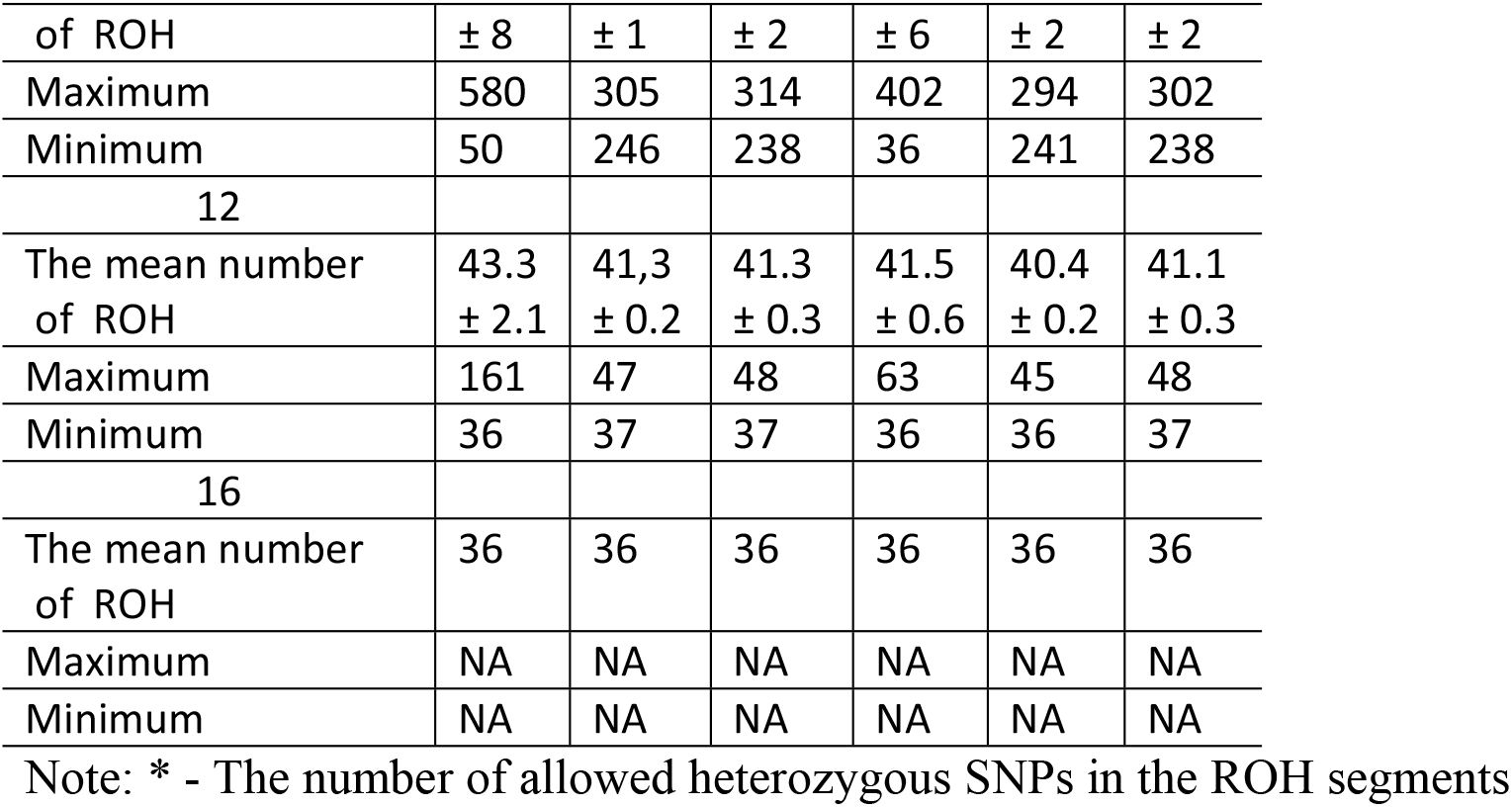
Estimated mean (± SE) ROH number in the herds based on 20 SNPs sliding runs

| Herd | 1 | 2 | 3 | 4 | 5 | 6 |
| --- | --- | --- | --- | --- | --- | --- |
| 0* |  |  |  |  |  |  |
| The mean number of ROH | 91<br>$\pm 6$ | 82<br>$\pm 1$ | 83<br>$\pm 1$ | 92<br>$\pm 14$ | 85<br>$\pm 1$ | 83<br>$\pm 1$ |
| Maximum | 336 | 194 | 138 | 731 | 106 | 111 |
| Minimum | 1 | 62 | 59 | 44 | 67 | 64 |
| 1 |  |  |  |  |  |  |
| The mean number of ROH | 157<br>$\pm 9$ | 142<br>$\pm 1$ | 143<br>$\pm 1$ | 146<br>$\pm 5$ | 147<br>$\pm 1$ | 145<br>$\pm 1$ |
| Maximum | 513 | 174 | 175 | 335 | 175 | 169 |
| Minimum | 5 | 117 | 114 | 82 | 118 | 120 |
| 2 |  |  |  |  |  |  |
| The mean number of ROH | 268<br>$\pm 10$ | 255<br>$\pm 1$ | 254<br>$\pm 2$ | 256<br>$\pm 4$ | 255<br>$\pm 2$ | 257<br>$\pm 2$ |
| Maximum | 631 | 284 | 287 | 294 | 285 | 288 |
| Minimum | 21 | 225 | 220 | 109 | 237 | 222 |
| 4 |  |  |  |  |  |  |
| The mean number of ROH | 524<br>$\pm 8$ | 535<br>$\pm 2$ | 540<br>$\pm 2$ | 533<br>$\pm 12$ | 534<br>$\pm 2$ | 542<br>$\pm 3$ |
| Maximum | 576 | 578 | 586 | 608 | 579 | 590 |
| Minimum | 133 | 494 | 484 | 39 | 492 | 502 |
| 8 |  |  |  |  |  |  |
| The mean number | 266 | 271 | 272 | 264 | 270 | 270 |
| of ROH | $\pm 8$ | $\pm 1$ | $\pm 2$ | $\pm 6$ | $\pm 2$ | $\pm 2$ |
| Maximum | 580 | 305 | 314 | 402 | 294 | 302 |
| Minimum | 50 | 246 | 238 | 36 | 241 | 238 |
| 12 |  |  |  |  |  |  |
| The mean number of ROH | 43.3<br>$\pm 2.1$ | 41.3<br>$\pm 0.2$ | 41.3<br>$\pm 0.3$ | 41.5<br>$\pm 0.6$ | 40.4<br>$\pm 0.2$ | 41.1<br>$\pm 0.3$ |
| Maximum | 161 | 47 | 48 | 63 | 45 | 48 |
| Minimum | 36 | 37 | 37 | 36 | 36 | 37 |
| 16 |  |  |  |  |  |  |
| The mean number of ROH | 36 | 36 | 36 | 36 | 36 | 36 |
| Maximum | NA | NA | NA | NA | NA | NA |
| Minimum | NA | NA | NA | NA | NA | NA |

Note: \* - The number of allowed heterozygous SNPs in the ROH segments

To have a comparable result to consecutive runs, 20 SNPs window was used further. Descriptive statistic for 20 SNPs sliding data is given in Table 3. Between herds mean number of ROH was insignificant for disallowed heterozygous SNPs in ROH. But, similar to consecutive runs after exclusion the most deviated cow (731 ROH segments) among fourth herd, the mean number ROH became 77.6 ± 2.0 and fourth herd significantly differed from third, fifth and sixth herds (P < 0.02) using t-test estimated with Bonferroni correction. The mean value increased six times from 86.0 ± 2.6 to 534.7 ± 2.5 when four heterozygous SNPs in ROH were allowed, then decreased to 36 when the number of heterozygous SNPs increased to sixteen.

The length of ROH segments were classified into the same five categories (1-2 Mb, 2-4 Mb, 4-8 Mb, 8-16 Mb, and >16 Mb) as it was done for consecutive runs (Table 4). The most abundant in the number of ROH was 1-2 Mb class as well. This fact indicates that with an increase in the number of allowed heterozygous SNPs, initially the number of short 1-2 Mb ROH had a bursting increase, and only then the number of longer ROH increased (Fig 2). Hence, close location of numerous shorter than 1 Mb ROH segments were identified in cow genome. Like this, starting from two allowed heterozygous SNPs, 4-8 Mb ROH segments began to increase, and starting from four heterozygous SNPs, 8-16 Mb and >16 Mb ROH segments likewise began to increase. Finally, only beyond eight heterozygous SNPs the number of 8-16 Mb ROH segments began to decrease. No doubt, sliding approach led to faster filling in of the cow genome with ROH segments compared to consecutive runs. It is necessary to pay attention to a sizable increase in the average length of ROH segments in the >16 Mb class when the number of allowed heterozygous SNPs exceeded 12 (S5 Table).

**Table 4.** Estimated mean (± SE) across herds ROH number in length classes based on 20 SNPs sliding runs

| Classes ( Mb) | 0-2 | 2-4 | 4-8 | 6-16 | >16 |
| --- | --- | --- | --- | --- | --- |
| 0* | 3190<br>± 241 | 1048<br>± 83 | 643<br>± 61 | 309<br>± 32 | 111<br>± 16 |
| 1 | 6488<br>± 554 | 1419<br>± 122 | 690<br>± 66 | 358<br>± 31 | 139<br>± 15 |
| 2 | 12205<br>± 1100 | 2444<br>± 220 | 817<br>± 88 | 385<br>± 39 | 152<br>± 14 |
| 4 | 22024<br>± 2139 | 8342<br>± 738 | 2169<br>± 208 | 545<br>± 54 | 185<br>± 19 |
| 8 | 2744<br>± 267 | 3333<br>± 312 | 4421<br>± 447 | 3795<br>± 379 | 2476<br>± 226 |
| 12 | 23<br>± 3 | 21.7<br>± 4.5 | 281<br>± 24 | 181<br>± 19 | 2071<br>± 193 |
| 16 | | | 249<br>± 23 | 124<br>± 12 | 1866<br>± 174 |

Note: \*- The number of allowed heterozygous SNPs in ROH

**Figure 2.**
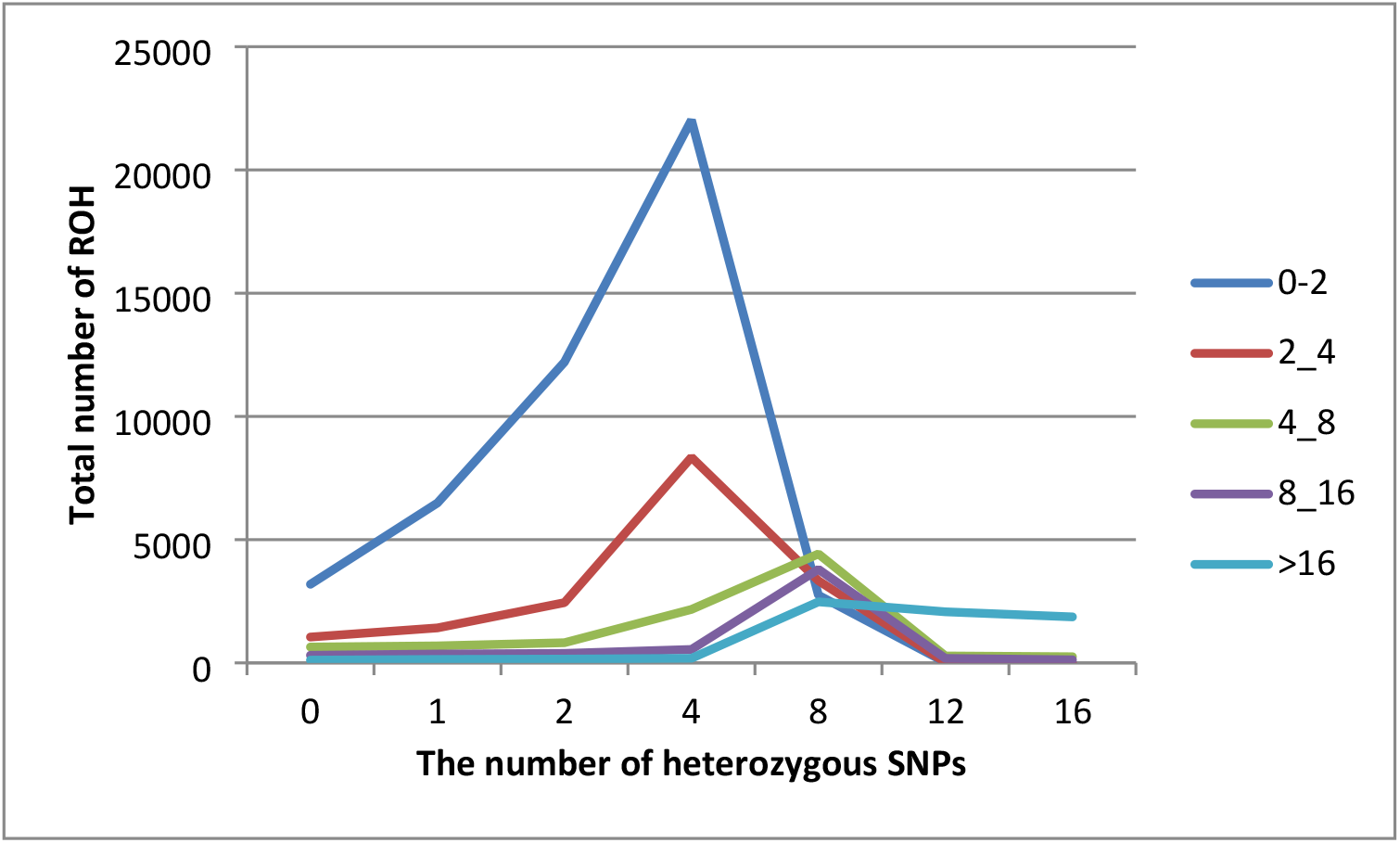
Distribution of ROH in five length classes based on sliding runs

#### ROH identification based on Plink tool

Plink software is widely used for ROH identification. Therefore, it was applied to estimate a possible data bias due to different software tools being used. It turn out, that the mean number of ROH obtained with Plink was 74.9 ± 1.9, which was significantly less of those calculated with detectRUNS based on sliding runs (P < 0.001). The fact, that Plink identified less ROH segments mainly in 1-2 Mb class than detectRUNS led to a larger mean ROH length in this class (S6 Table *vs*. S3 Table and S5 Table). Thus, the data obtained for the shortest ROH length class is highly dependent on the software and parameters used.

#### Comparative analysis of consecutive and sliding runs

Comparative analysis of the consecutive and sliding data led to the following conclusions. If heterozygous SNPs were disallowed the consecutive runs identified slightly more ROH than sliding runs (94.4 ± 2.7 *vs*. 86.0 ± 2.8, P < 0.05). The shorter SNPs window at sliding runs or minimal number of SNPs needed to define segment at consecutive runs were, the more abundant in the ROH number was 1-2 Mb class (Table 2 and Table 4 *vs*. S2 Table and S6 Table). If the number of allowed heterozygous SNPs in ROHs continued to increase, the distribution of ROH segments peaked faster (on the number of heterozygous SNPs scale) when the sliding runs were used (Fig 1 *vs*. Fig 2). Necessary to note that the distributions based on consecutive runs had several ROH maximum (Fig 1) while those based on sliding runs had only two ROH maximum (Fig 2). There are further points to be made here. The cow genome had been filled in with ROH segments when sixteen heterozygous SNPs were allowed at sliding runs while their number was increased up to 240 for consecutive runs (Fig 3). Further, as it is shown on Fig 3, the sliding method identified the less number of ROH segments with smaller number of allowed heterozygous SNPs. Summarizing comparative analysis of both methods, we can conclude the large differences between findings of these methods. In the section below will be discussed how the ROH segments are distributed in cow genome.

**Figure 3.**
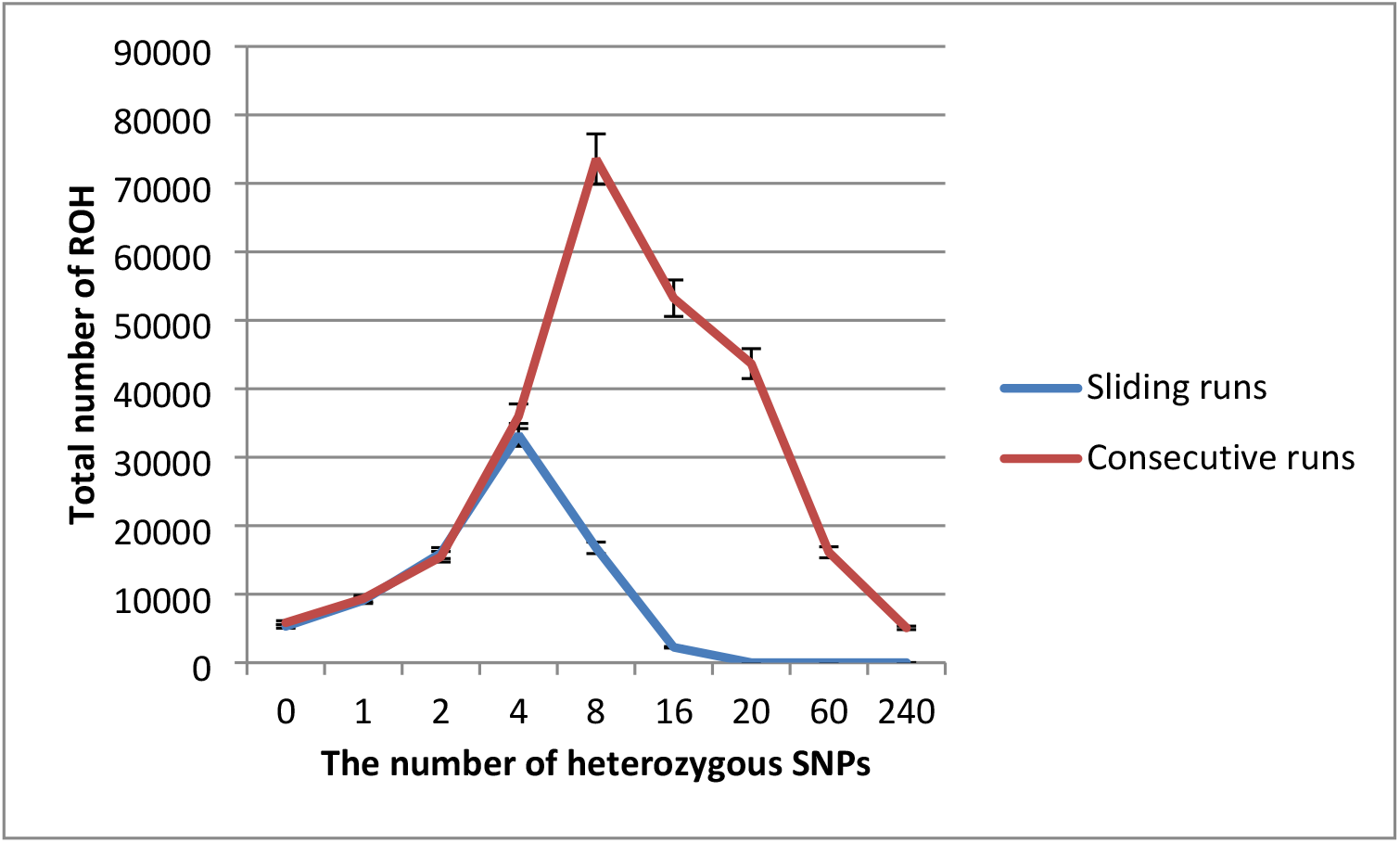
Total distribution of ROH in the cow genome

#### Evaluation of the distances between ROH segments

The number of allowed heterozygous SNPs might be converted into nucleotide distance. For example, the distances between two non-interrupted by heterozygous SNPs ROH segments (**niROH**) separated by one heterozygous SNP will be equal to the mean distance between SNPs in the cow genome multiplied by two. If two heterozygous SNP were allowed then, either the distances between three niROH would be the same as for one heterozygous SNP or it would be double of mean length between SNPs in genome, plus yet one mean distance, if two niROH were separated by two heterozygous SNPs. Anyway, if the number of allowed heterozygous SNPs was being continued to increase, the mean distance between niROH also was increasing. That is the number of allowed heterozygous SNPs may be used as a measure of the distance between niROH. Summarizing the above mentioned reasoning, the upper limit of distance between niROH segments will be D = (n + 1)m, where n is the number of allowed heterozygous SNPs and m is the mean distance between SNPs in cow genome equal to 52.1 Kb (2508.706681 Mb (genome length covered by SNPs) / 48108 (number of genotyped SNPs) = 52.1 Kb). Based on this formula it is possible to provide some estimates for each length class about maximum and minimum distance between niROH and their length in ROH clusters. For each class there was a threshold beyond of which being impossible to form a ROH clusters from niROH that would not exceed the maximum length for this class. Then, a longer length class will begin to form. But, SNPs unevenly located in the cow genome [26]. Owing to this phenomenon and different length of niROHs, the number of heterozygous SNPs suitable for every class length limit would be hardly to predict if not based on the mean distances between SNPs.

#### Calculation of the distances between ROH segments

Let us calculate upper and lower limits between niROH in the length classes. If one heterozygous SNP was allowed then mean distance between two niROH segments would be 104 Kb and those niROH form an initial ROH clusters. If eight heterozygous SNPs were allowed, then different number of niROH segments might formed the ROH clusters belonging to the 1-2 Mb class. In this case, the maximum mean upper limit on distance between two niROH will be 450 Kb. Otherwise, this distance should begin to decrease if the number of niROH will increase. Given that maximum length 2 Mb could not be exceeded for 1-2 Mb class, then minimal mean lower limit on distances between niROH would be nine 133 Kb length niROH separated by eight 100 Kb heterozygous SNP segments. Based on Fig 1 it can be concluded, with an increase in the number of heterozygous SNPs over 8, the number of niROH clusters satisfying 1–2 Mb class began to decrease. Further increase in the number of heterozygous SNPs led to an increase in the number of ROH in the next class. The same calculation is valid for other length classes but each of them has its own maximal and minimal number of niROH and distances between them.

Since the distances between SNPs in the cattle genome vary considerably [27], some SNPs may be located extremely close to each other. It can be illustrated for 1-2 Mb class when more than 60 heterozygous SNPs were allowed (Table 2). Indeed, for 60 allowed heterozygous SNPs in ROH, summed mean distance between 60 niROH would be ≈ 3.5 Mb, that exceeds the length 2 Mb for 1–2 Mb class. As it was mentioned above there were noticeable differences in results of two methods. The sliding window method underestimated the distances between ROH and, therefore, the position of the peaks were shifted and merged relative to those for consecutive runs (Fig 1 *vs*. Fig 2).Therefore, for these methods the number of ROH and distances between niROH considerably differed and obtained findings were inconsistent. Visualization of ROH formation owing to niROH clusters is illustrated on Fig S1 – S3, on which the BTA 28 was chosen having the smallest number of ROH segments. When heterozygous SNPs were disallowed, only niROH were identified on BTA 28 (Fig S1). These niROH segments 1-18 Mb length scattered along BTA 28. When one heterozygous SNP was allowed, additional niROH clasters forming new short ROH segments can be seen on BTA 28 (Fig S2). Their length was mostly 1-2 Mb. As the number of heterozygous SNPs increased to two, new niROH clusters forming ROH segments different length can be seen on the chromosome 28 (Fig S3).

#### Distribution of ROH in the cow genome

In order to evaluate the chromosomes with largest number of ROH segments, calculation of ROH proportion in them based on normalization to each chromosome length was performed. For both runs the ranked chromosomes list is given in Table 5. Of 29 chromosomes, the highest proportion of ROHs was calculated at BTA14, BTA16, and BTA7 not BTA1 (seventh position in the list), BTA2 (22th position in the list) and BTA3 (15th position in the list). Thus, the number of ROH in chromosomes was not directly proportional to the chromosome length. Spearman’s correlation between consecutive and sliding runs data set was r = 1.0 (P ≤ 2.0·10^−7^).

**Table 5.** Normalized proportion of ROH in the cow chromosomes

|  |  |  |  |  |  |  |  |  |  |  |
| --- | --- | --- | --- | --- | --- | --- | --- | --- | --- | --- |
| Chromosome | 14 | 16 | 7 | 26 | 8 | 13 | 1 | 22 | 17 | 4 |
| Consecutive runs* | 1.350 | 1.280 | 1.240 | 1.200 | 1.190 | 1.120 | 1.110 | 1.098 | 1.060 | 1.050 |
| Sliding runs | 1.388 | 1.295 | 1.218 | 1.170 | 1.169 | 1.161 | 1.086 | 1.011 | 1.050 | 1.030 |
| Chromosome | 20 | 6 | 19 | 21 | 3 | 24 | 10 | 12 | 11 | 5 |
| Consecutive runs | 1.024 | 1.018 | 1.016 | 0.964 | 0.955 | 0.929 | 0.928 | 0.926 | 0.920 | 0.911 |
| Sliding runs | 0.980 | 0.999 | 1.010 | 0.965 | 0.963 | 0.955 | 0.931 | 0.918 | 0.909 | 0.911 |
| Chromosome | 9 | 2 | 25 | 29 | 23 | 15 | 18 | 27 | 28 |  |
| Consecutive runs | 0.908 | 0.900 | 0.876 | 0.844 | 0.814 | 0.782 | 0.754 | 0.743 | 0.679 |  |
| Sliding runs | 0.902 | 0.898 | 0.920 | 0.865 | 0.851 | 0.800 | 0.743 | 0.774 | 0.680 |  |

Note: \* - The values of proportions were ranged from maximum to minimum only for consecutive runs

As can be seen on Fig 1 and Fig 2, for both methods the distributions of ROH number from allowed number of the heterozygous SNPs within them was similar to normal. To measure a proximity to the normal distribution Q-Q testing was used. It was shown their similarity to normal distribution but to varying degree for different length classes (S4 Fig –S13 Fig). It should be borne in mind that for small samples a greater variation is expected. Obviously, the distributions for classes 1-2 Mb, 2-4 Mb and 8-16 Mb had a positive skewness and light tailed form, while for classes 8-16 Mb and >16 Mb the distributions were the most similar to normal except class >16 Mb for sliding runs (S13 Fig).

#### Inbreeding

To estimate a level of inbreeding in the herds, the mean inbreeding coefficient across herds was calculated (Table 6 and Table 7). If heterozygous SNPs were disallowed, then the mean inbreeding coefficients amounted to 0.111 ± 0.003 and 0.104 ± 0.003 for consecutive and sliding runs respectively, and difference between them was insignificant. The FROH estimated with Plink was 0.105 ± 0.004 what consistent with that for sliding runs. The largest variability of inbreeding was observed in fourth herd. This was mainly due to highly inbred cow in this herd. Exclusion this cow led to F_ROH_ 0.096 ± 0.005 and 0.089 ± 0.005 for consecutive and sliding runs. In this case, fourth herd significantly differed from herds 1 and herd 5 (P < 0.01) and herd 2 (P < 0.02), given t-test estimated with Bonferroni correction. It should be noted that in this herd insemination of the cows only from The Netherland bulls has occurred, while for other herds the bulls semen from North America and Canada were used as well. After excluding the highly inbreed cow from fourth herd, the average F_ROH_ in this herd became less than for other herds. This result indicates the correct selection of the bulls, even they were used from a single country. It was shown, the forth herd largely deviated from other herds when it was measured by fixation index and Principal Component Analysis [32]. When one heterozygous SNP was allowed in ROH, then F_ROH_ were 0.145 ± 0.003 and 144 ± 0.003 based on consecutive and sliding runs. Thus, an allowing of even one heterozygous SNP led to significant increase in F_ROH_ (P ≤ 0.001). Therefore, to evaluate inbreeding in the herds, the heterozygous SNPs should not be allowed in ROH due to sizable bias. Distribution of F_ROH_ from the number of heterozygous SNPs is shown on Figure 4. Initially, a relatively gradual increase of F_ROH_ occurred. Then, an increasing greatly accelerated. Finally, increasing gradually slowed down again. In fact, this curve indicated the relative distances between niROH on heterozygous SNPs scale. From Fig 4 it can be seen that 20% of niROH were located at short nucleotide distances less 1 Mb, 80% of ROH clusters formed from niROH were located at intermediate distances, and at least 20% of niROH clusters were located at most longest distances. Thus, in the cow genome there is a complex hierarchical structure of niROH clusters.

**Table 6.** Estimated mean (± SE) inbreeding coefficient in the herds based on 20 SNPs consecutive runs

| Herd | 1 | 2 | 3 | 4 | 5 | 6 |
| --- | --- | --- | --- | --- | --- | --- |
| 0* |  |  |  |  |  |  |
| Inbreeding coefficient | 0.117<br>± 0.004 | 0.112<br>± 0.002 | 0.105<br>± 0.002 | 0.111<br>± 0.016 | 0.116<br>± 0.003 | 0.105<br>± 0.003 |
| Maximum | 0.0006 | 0.060 | 0.068 | 0.047 | 0.075 | 0.071 |
| Minimum | 0.227 | 0.153 | 0.160 | 0.779 | 0.166 | 0.158 |
| 1 |  |  |  |  |  |  |
| Inbreeding coefficient | 0.153<br>± 0.006 | 0.143<br>± 0.002 | 0.143<br>± 0.002 | 0.147<br>± 0.018 | 0.147<br>± 0.002 | 0.137<br>± 0.003 |
| Maximum | 0.003 | 0.091 | 0.091 | 0.072 | 0.104 | 0.102 |
| Minimum | 0.367 | 0.184 | 0.184 | 0.908 | 0.191 | 0.192 |
| 2 |  |  |  |  |  |  |
| Inbreeding coefficient | 0.208<br>± 0.008 | 0.195<br>± 0.002 | 0.189<br>± 0.002 | 0.198<br>± 0.017 | 0.199<br>± 0.002 | 0.190<br>± 0.003 |
| Maximum | 0.010 | 0.139 | 0.148 | 0.102 | 0.158 | 0.155 |
| Minimum | 0.511 | 0.235 | 0.229 | 0.948 | 0.240 | 0.246 |
| 4 |  |  |  |  |  |  |
| Inbreeding coefficient | 0.384<br>± 0.011 | 0.372<br>± 0.002 | 0.369<br>± 0.002 | 0.377<br>± 0.01 | 0.376<br>± 0.002 | 0.369<br>± 0.002 |
| Maximum | 0.053 | 0.317 | 0.333 | 0.229 | 0.349 | 0.333 |
| Minimum | 0.742 | 0.407 | 0.406 | 0.972 | 0.416 | 0.410 |
| 8 |  |  |  |  |  |  |
| Inbreeding coefficient | 0.787<br>± 0.008 | 0.788<br>± 0.001 | 0.788<br>± 0.001 | 0.791<br>± 0.005 | 0.790<br>± 0.001 | 0.788<br>± 0.001 |
| Maximum | 0.396 | 0.763 | 0.764 | 0.667 | 0.773 | 0.772 |
| Minimum | 0.934 | 0.809 | 0.813 | 0.981 | 0.814 | 0.808 |
| 16 |  |  |  |  |  |  |
| Inbreeding coefficient | 0.9399<br>± 0.0009 | 0.9395<br>± 0.0002 | 0.9391<br>± 0.0002 | 0.9399<br>± 0.001 | 0.9397<br>± 0.0002 | 0.9391<br>± 0.0002 |
| Maximum | 0.903 | 0.934 | 0.935 | 0.924 | 0.936 | 0.936 |
| Minimum | 0.964 | 0.944 | 0.944 | 0.985 | 0.943 | 0.943 |
| 20 |  |  |  |  |  |  |
| Inbreeding coefficient | 0.9492<br>± 0.0008 | 0.9488<br>± 0.0002 | 0.9487<br>± 0.0002 | 0.9495<br>± 0.0009 | 0.9489<br>± 0.0002 | 0.9488<br>± 0.0002 |
| Maximum | 0.919 | 0.945 | 0.945 | 0.938 | 0.946 | 0.945726 |
| Minimum | 0.971 | 0.953 | 0.952 | 0.987 | 0.953 | 0.952828 |
| 100 |  |  |  |  |  |  |
| Inbreeding coefficient | 0.9814<br>± 0.0002 | 0.98124<br>± 0.00009 | 0.98117<br>± 0.00009 | 0.9814<br>± 0.0002 | 0.98115<br>± 0.00008 | 0.9814<br>± 0.0001 |
| Maximum | 0.975 | 0.979 | 0.979 | 0.978 | 0.980 | 0.979 |
| Minimum | 0.986 | 0.983 | 0.982 | 0.990 | 0.982 | 0.983 |
| 240 |  |  |  |  |  |  |
| Inbreeding coefficient | 0.9872<br>± 0.0001 | 0.98641<br>± 0.00007 | 0.98635<br>± 0.00007 | 0.9864<br>± 0.0001 | 0.98626<br>± 0.00007 | 0.98642<br>± 0.00007 |
| Maximum | 0.983 | 0.984 | 0.985 | 0.985 | 0.985 | 0.985 |
| Minimum | 0.989 | 0.988 | 0.987 | 0.990 | 0.988 | 0.988 |

Note: \* - The number of allowed heterozygous SNPs in ROH

**Table 7.** Estimated mean (± SE) inbreeding coefficient in the herds based on 20 SNPs sliding runs

| Herd | 1 | 2 | 3 | 4 | 5 | 6 |
| --- | --- | --- | --- | --- | --- | --- |
| Zero heterozygous SNP in ROH |  |  |  |  |  |  |
| Inbreeding coefficient | 0.110<br>± 0.004 | 0.105<br>± 0.002 | 0.098<br>± 0.002 | 0.103<br>± 0.015 | 0.109<br>± 0.003 | 0.098<br>± 0.003 |
| Minimum | 0.0003 | 0.055 | 0.063 | 0.040 | 0.070 | 0.065 |
| Maximum | 0.204 | 0.146 | 0.153 | 0.739 | 0.158 | 0.151 |
| One heterozygous SNP in ROH |  |  |  |  |  |  |
| Inbreeding coefficient | 0.154<br>± 0.007 | 0.143<br>± 0.002 | 0.137<br>± 0.002 | 0.146<br>± 0.018 | 0.148<br>± 0.002 | 0.138<br>± 0.003 |
| Minimum | 0.002 | 0.092 | 0.094 | 0.071 | 0.104 | 0.104 |
| Maximum | 0.392 | 0.186 | 0.184 | 0.938 | 0.192 | 0.194 |
| Two heterozygous SNP in ROH |  |  |  |  |  |  |
| Inbreeding coefficient | 0.227<br>± 0.01 | 0.213<br>± 0.002 | 0.207<br>± 0.002 | 0.217<br>± 0.018 | 0.217<br>± 0.002 | 0.208<br>± 0.003 |
| Minimum | 0.010 | 0.157 | 0.167 | 0.108 | 0.173 | 0.173 |
| Maximum | 0.583 | 0.254 | 0.244 | 0.980 | 0.254 | 0.258 |
| Four heterozygous SNP in ROH |  |  |  |  |  |  |
| Inbreeding coefficient | 0.474<br>± 0.012 | 0.463<br>± 0.002 | 0.461<br>± 0.002 | 0.470<br>± 0.012 | 0.467<br>± 0.002 | 0.463<br>± 0.002 |
| Minimum | 0.072 | 0.417 | 0.417 | 0.298 | 0.439 | 0.429 |
| Maximum | 0.859 | 0.503 | 0.504 | 0.989 | 0.502 | 0.506 |
| Eight heterozygous SNP in ROH |  |  |  |  |  |  |
| Inbreeding | 0.924 | 0.929 | 0.928 | 0.930 | 0.930 | 0.929 |
| coefficient | $\pm 0.007$ | $\pm 0.0005$ | $\pm 0.0007$ | $\pm 0.003$ | $\pm 0.0006$ | $\pm 0.0007$ |
| Minimum | 0.562 | 0.918 | 0.915 | 0.826 | 0.921 | 0.919 |
| Maximum | 0.988 | 0.940 | 0.941 | 0.100 | 0.942 | 0.942 |
| Twelve heterozygous SNP in ROH |  |  |  |  |  |  |
| Inbreeding coefficient | 0.988<br>$\pm 0.0005$ | 0.989<br>$\pm 4.6 \cdot 10^{-5}$ | 0.989<br>$\pm 6.15 \cdot 10^{-5}$ | 0.989<br>$\pm 0.0001$ | 0.989<br>$\pm 6.16 \cdot 10^{-5}$ | 0.989<br>$\pm 5.58 \cdot 10^{-5}$ |
| Minimum | 0.961 | 0.987 | 0.988 | 0.985 | 0.987 | 0.988 |
| Maximum | 0.9898 | 0.9897 | 0.9897 | 0.9898 | 0.9898 | 0.9896 |
| Sixteen heterozygous SNP in ROH |  |  |  |  |  |  |
| Inbreeding coefficient | 0.989824 | 0.989824 | 0.989824 | 0.989824 | 0.989824 | 0.989824 |
| Minimum | NA | NA | NA | NA | NA | NA |
| Maximum | NA | NA | NA | NA | NA | NA |

**Figure 4.**
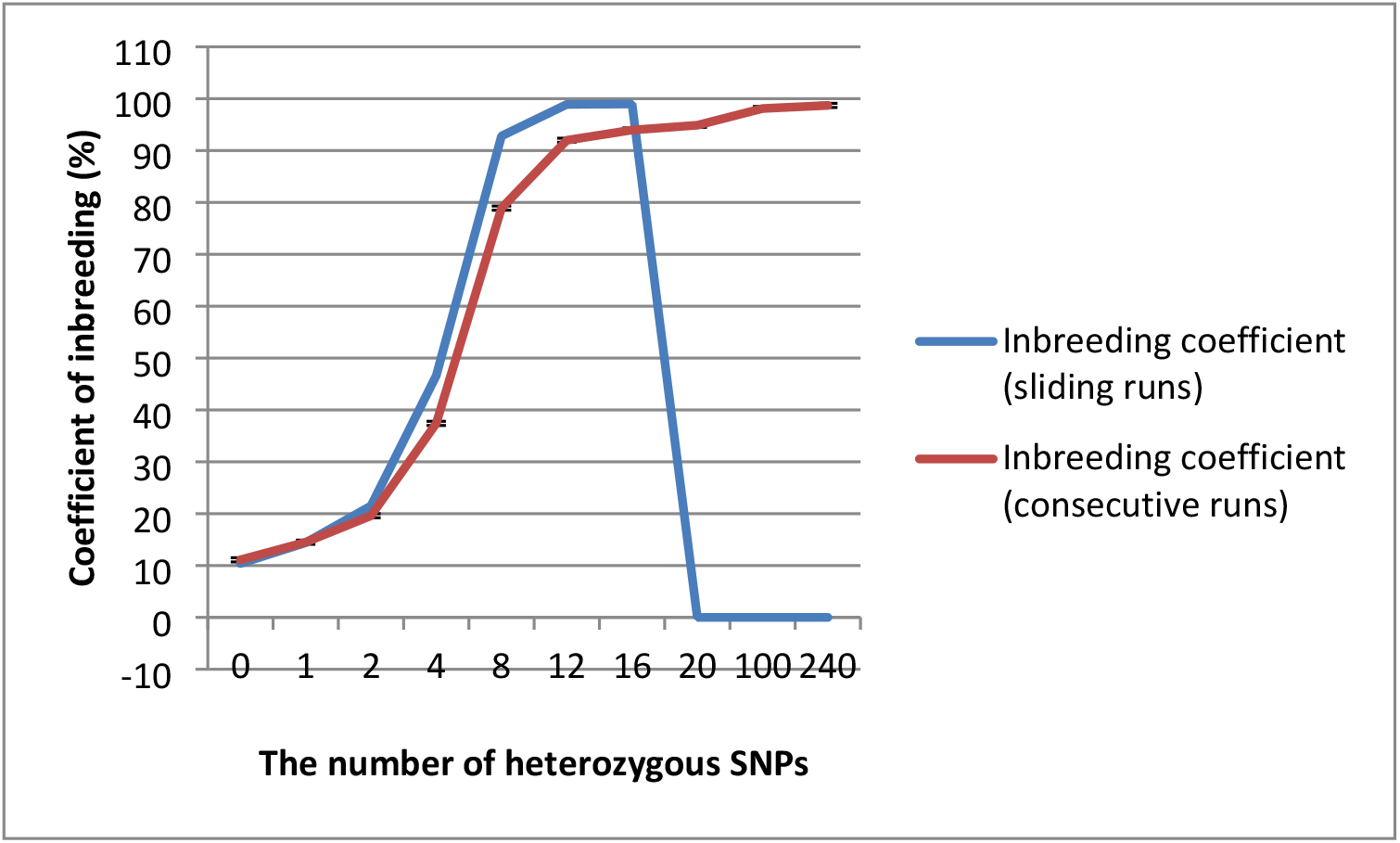
Dependence of the inbreeding coefficient on the number of heterozygous SNPs

## Discussion

Over past decade the ROH approach has been widely used both on humans [3] and farm animals [2]. Hallmark of ROH studies is a lot of software and thresholds criteria to be used in them. The most widely applied software tools to identify ROH stretches are either sliding window or consecutive runs. We preferred detectRUNS where both approach were implemented [9]. The consecutive runs resulted in the average number of ROH 94.4 ± 2.7 while sliding runs 86.0 ± 2.6. This values for North American [8], Italian [16], European Holstein [21], and Polish Holstein Black-and White variety [28] are 82.3 ± 9.8(SD), 81.7 ± 9.7(SD), 74.6 ± 2.3(SE), and 53.3 ± 7.3(SD) respectively. The first three values do not differ significantly from our ones, while the value for Polish cattle considerably differs. It was shown that allowance of even one heterozygous SNP in ROH significantly increases the number of ROH by 56.9 and 60.7 points for consecutive and sliding runs respectively (Table 1 and Table 3). A limited number of studies investigated the effect of allowed heterozygous SNPs on ROH data. Howrigan et al. [13] recommended disallowing any heterozygous SNPs in ROH, while Ferencakovic et al. [22] suggested the number of heterozygous SNPs allowed should be determined separately for each ROH length of interest and for each SNPs density.

The relative frequency of ROH number in different length classes for consecutive runs were 61.4% (1–2 Mb), 19.8% (2-4 Mb), 11.3% (4–8 Mb), 5.5% (8–16 Mb) and (longer than 16 Mb) 1.9%, while for sliding runs these values were 60%, 19.8%, 12.1%, 5.8%, 2.1%. Thus, the largest number of ROH was identified in the shortest 1–2 Mb class. By Plink-running of the cow genome was identified the following frequency of ROH in five categories 52% (0-2 Mb), 25% (2–4 Mb), 14% (4–8 Mb), 7% (8-16 Mb) and (longer than 16 Mb) 2.5% whose distribution is slightly different from those identified with detectRUNS. For North American Holstein animals these values were 43.5%, 23.9%, 17.7%, 10.5%, and 4.7% [8]. Corresponding values for Italian Holstein bulls were 56.9%, 20.8, 11.9, 7.2, and 3.7% [16] and Polish Holstein 21%, 37.1%, 23.1%, 13.3, and 5.6% [28]. Thus, when we used detectRUNS for genome scanning of our local Holsteinzied cows, we obtained an abundant number of short ROH resulted from haplotypes reflecting the ancient relationship within breeding animals. But, when we used Plink, the values were similar to those for American and Italian data. It should be considered that authors of the article [28] used cgaTOH software and their data considerably differ from other data. Whether this result was due to cgaTOH software or/and breeding management requires furthers analysis. Estimation of the true amount of short ROH is important, since 0.1–3 Mb ROH segments have the more number of deleterious variants than segments greater of 3 Mb [20]. Furthermore, to evaluate genomic estimated breeding value (GEBV), the short ROH is important for genomic ROH-based relationship matrix (GROH) construction [29]. According to our estimates, the most number of ROH was in 1–2 Mb class, which included niROHs shorter than 1 Mb, located at short distances and forming 1-2 Mb length ROH clusters, thereby determining the structure of the cattle genome (Fig S1-S3). Finally, the results of the animal ROH genome scanning can substantially depend on not only from selection strategy, but software used. This fact should be taken into account in comparative analysis of the ROH data.

Currently evaluated with detectRUNS the mean inbreeding coefficient for six herds was 0.111 ± 0.003 and 0.104 ± 0.003. It is equal to 0.105 ± 0.004 based on Plink runs. These values do not differ significantly (t-test). But, F_ROH_ increased significantly (P < 0.001) from 0.111 ± 0.003 to 0.145 ± 0.003, when one heterozygous SNP was allowed in ROH (Table 6). However, Mastrangelo et al. [24] showed that the differences between F_ROH_ estimated using one, two, and three heterozygous SNPs were very small in Italian Holstein (0.042, 0.045, 0.049) when F_ROH_ was estimated for >4 Mb ROH. F_ROH_ for American Holstein was 0.12 in 2011 year and after applying genomic selection it increased until 0.15 in 2018 year [8]. For European [21], Italian [16], and Polish Holstein [26] these values were 0.108 ± 0.006 (SE), 0.116 ± 0.001 (SE), and 0.118 ± 0.027 (SD) respectively. In the studies cited above, ROH data were based on 50k arrays, thereby not shorter than 1 Mb ROH segments were identified. Thus, short ROH were underestimated that might be led to an inaccurate estimate of inbreeding occurred several tenth years ago. The accurate knowledge of inbreeding in the herds is necessary for both calculation of inbreeding trend and evaluation of the selection strategies. In short ROH the deleterious mutations may be concentrated [20]. To overcome this problem a high density arrays or whole genome sequencing (WGS) should be used. Comparing 50K and HD panels provide evidence that data from 50K panel lead to imprecise determination of short ROH segments [22]. However, detection the ROH based on high-density or 50k chip data was shown to be able to give the estimates of current inbreeding the most similar to ROH values obtained from sequence data [30]. Bhali et al. [31] provided the comprehensive WGS data for Branvich cattle. The medium – sized ROH (0.1–2 Mb) were the most frequent class (50.46%) and contributed most (75%) of the total genomic inbreeding, while short ROH 50 – 100 Kb occurred almost as frequent (49.17%) as medium-sized ROH, they contributed only 19.52% to total genomic inbreeding. These findings provide an accurate estimate of short ROH in the cattle genome and their input to total inbreeding. The average F_ROH_ estimated from WGS data was 0.14 in Branvich cattle. This value is less than WGS F_ROH_ in Holstein 0.18 [31]. Thus, the results inferred from WGS provide a larger F_ROH_ than those from SNPs array (0.15 *vs*. 0.18). Thus, 50k panel can not accurately capture inbreeding occurred several tenth years in the past.

It may be assumed that for a herd or population has to be an event horizon beyond, which it is impossible to obtain only from ROH data valid information about their breeding history events. Our local population with both reduced effective population size, ongoing admixture and inbreeding throughout its breeding history, accompanied by recombination, should has the greatest number of short ROH less than 1 Mb in herds. These short ROH can be considered as a relict ROH segments formed due to some population events, such as admixture, drift, bottleneck, and inbreeding occurred many decades ago. Previously, with Principal Component Analysis the bottleneck in our local herds was not proved [32]. An accurate interpretation of these ROH can be troublesome without knowledge of the herd management history. Nevertheless, it is very important to know the true number of short ROHs at the animals analyzed (see above mentioned WGS data). Thus, the event horizon might be depended as from pedigree information, ROH length profile (used SNPs array or WGS) as well as on algorithm-defined approach for ROHs identification. However, ROH segments shorter than 1 Mb can be considered as being beyond the event horizon due to significant shortening by recombination. An addition, short ROH characterized by strong LD among markers are not always considered autozygous but, nevertheless, are due to mating to distantly related animals [33]. Summarizing, it should be assume that inbreeding data based on array can only be relatively correct for an ROH larger than 1 Mb (no more 50 generation back).

### Distribution of ROH in the cow genome

In several studies have noticed an uneven distribution of ROH in the chromosomes [e.g. 10, 14, 34]. Giving the number of ROH in the chromosomes, their proportion normalized on each chromosome length was calculated (Table 5). Of 29 chromosomes, the most fill in with ROH segments were BTA14, BTA16 and BTA7 for both runs approach used. Purfield at al. [17] noticed that the largest extent of overlapping ROH positions among studied breeds had ROH segments on BTA14 and BTA16. The genome regions with the highest frequency of ROH occurrence were called “ROH islands” [35, 36]. The ROH islands on BTA14 and BTA16 were identified among Polish Holstein-Friesian animals [26]. In American Holstein ROH distribution was more variable among the genomes of selected animals in comparison to a more even ROH distribution at unselected animals [37]. Further, in this study more than 40 genetic regions under selection on BTA2, BTA7, BTA16 and BTA20 were identified. Regions with high level of ROH for American and New Zealand Jersey cows and bulls were detected on BTA3 and BTA7 [10]. One strongest ROH region common for Kholmogor and Holstein breeds and one region common for Yaroslavl and Holstein breeds was found on BTA14 and BTA16 [21]. An extremely non-uniform ROH patterns among bovine populations of Angus, Brown Swiss, and Fleckvieh breeds were mainly located on BTA6, BTA7, BTA16, and BTA21 [34]. On BTA7 was found the highest number of ROH islads among all Neilore breed lineages [38]. Furthermore, on BTA14 in dairy Gyr breed (Bos Indicus) has shown an enrichment of genes affecting traits of interest for dairy breeds [39]. Goszczynski et al. [40] analyzed ROH *>*16 Mb (three generations since the common ancestor) in highly inbred Retinta bulls. Among other chromosomes the highest occurrence of ROH was found on BTA7. Summarizing studies cited above, it can be suggested that BTA7 is outstanding regarding ROH islands occurrence in the cattle genome but there is no overall direct relationship between frequency of ROH segments in chromosomes and identified ROH islands there.

Having determined proportion of ROHs in the chromosomes, it is necessary to define a type of distribution over which ROH segments located in genome. Considering the dependence of the ROH number from the number of allowed heterozygous SNPs in them, promoted to realize this task. For this, recalculation of the number of allowed heterozygous SNPs in ROH into distances between niROH was performed. To further elucidate the structure of ROH segments they were classified into five length classes. It turned out that in each class the number of ROH segments, which were formed from niROH were distributed similar to the normal distribution (Fig 1 and Fig 2). It was true for total set of ROH segments as well (Fig 3).This indicates a cluster structure of ROH in the cow genome. The ROH clusters form a hierarchical structure from short to longer as can be visualized on Fig S1-S3. As was shown, the number of short ROH was highly depended on the software and genotyping method used. Moreover, we show that the consecutive runs more accurately identified ROH pattern in cow genome. Nevertheless, it would like to point out that both methods coincide in estimate of ROH distribution type being similar to normal. Taken together our findings it should be admit about uneven distribution ROH segments in the cow genome as a result of different population events occurred during it history.

## Conclusions

Our analysis of the ROH data was able to estimate that consecutive runs the most accurately identified ROH in the cow genome. It was shown that missing SNPs had not sizable effect on the number of ROH, while an allowance of even one heterozygous SNP in ROH had significant effect. Therefore, caution should be taken to allow any number of heterozygous SNPs in ROH. The mean inbreeding coefficient for our local herds was 0.111 ± 0.003, which was not differ from those for the Holstein breed over the world. It was suggested that ROH segments had a tendency for clustering in the cow genome. Distribution of ROH clusters in the cow genome was similar to normal. Moreover, the number of ROH segments in the chromosomes did not depend on their length and the most abundant in ROH segments were BTA14, BTA7, and BTA18.

## Author contribution

## Supporting information

**S1 Table. Estimated mean (± SE) ROH number in the herds based on 15 SNPs consecutive runs.**

(DOCX)

**S2 Table. Estimated mean (± SE) across herds ROH number in length classes based on 15 SNPs consecutive runs.**

(DOCX)

**S3 Table. The mean (± SE) across herds ROH length in the classes based on 20 SNPs consecutive runs.**

(DOCX)

**S4 Table**. **Estimated mean (± SE) ROH number in the herds based on 15 SNPs sliding runs.**

(DOCX)

**S5 Table**. **The mean across herds ROH length in the classes based on 20 SNPs sliding runs**.

(DOCX)

**S6 Table**. **The mean (± SE) across herds ROH length in the classes based on Plink runs.**

(DOCX)

## Figures

**S1 Fig. Distribution of ROH segments along BTA 28 based on consecutive runs.**

Each row represents a cow from first herd and each column represents a genotyped SNPs position. Each ROH is a niROH. The heterozygous SNPs were disallowed.

(PDF)

**S2 Fig. Distribution of ROH segments along BTA 28 based on consecutive runs.**

Each row represents a cow from first herd and each column represents a genotyped SNPs position. Each ROH consists of some niROH number. One heterozygous SNPs were allowed.

(PDF)

**S3 Fig. Distribution of ROH segments along BTA 28 based on consecutive runs.**

Each row represents a cow from first herd and each column represents a genotyped SNPs position. Each ROH consists of some niROH number. Two heterozygous SNPs were allowed.

(PDF)

**S4 Fig. Q-Q plot based on consecutive runs**.

Distribution of ROH segments from 1-2 Mb class.

(PDF)

**S5 Fig. Q-Q plot based on consecutive runs**.

Distribution of ROH segments from 2-4 Mb class.

(PDF)

**S6 Fig. Q-Q plot based on consecutive runs**.

Distribution of ROH segments from 4-8 Mb class.

(PDF)

**S7 Fig. Q-Q plot based on consecutive runs**.

Distribution of ROH segments from 8-16 Mb class.

(PDF)

**S8 Fig. Q-Q plot based on consecutive runs.**

Distribution of ROH segments from >16 Mb class.

(PDF)

**S9 Fig. Q-Q plot based on sliding runs**.

Distribution of ROH segments from 1-2 Mb class.

(PDF)

**S10 Fig. Q-Q plot based on sliding runs**.

Distribution of ROH segments from 2-4 Mb class.

(PDF)

**S11 Fig. Q-Q plot based on sliding runs**.

Distribution of ROH segments from 4-8 Mb class.

(PDF)

**S12 Fig. Q-Q plot based on sliding runs**.

Distribution of ROH segments from 8-16 Mb class.

(PDF)

**S13 Fig. Q-Q plot based on sliding runs**.

Distribution of ROH segments from >16 Mb class.

**(PDF)**

